# Antioxidant, Anti-inflammatory and Anticancer activities of Methanolic Extract of *Thottea siliquosa:* An In Vitro study

**DOI:** 10.1101/2020.03.29.014324

**Authors:** Amrutha Koottasseri, Amal Babu, Anna Augustin, Joice Tom Job, Arunaksharan Narayanankutty

## Abstract

**Background and Aim:** Oxidative stress and inflammation are the predominant cause of chronic diseases including cancers, thereby making it as an essential target for preventing these disorders. Various natural products and plants have been shown to prevent the process of free radical induced damages. The present study evaluated the biological properties of *Thottea siliquosa,* belonging to the family Aristolochiaceae, which is a traditionally used Ayurvedic plant.

**Experimental Procedure:** Antioxidant assays carried out were DPPH, FRAP, hydrogen peroxide scavenging and hemolysis inhibition assay; nitric oxide and lipoxygenase inhibition assays were used for anti-inflammatory studies. Anticancer activity was done using human endometrial and breast cancer cells by MTT assay.

**Results and Conclusion:** The results of the present study showed antioxidant properties for the methanolic crude extract of *T. siliquosa* as DPPH radical scavenging (110.40± 4.5μg/mL), FRAP capacity (41.1± 6.2), peroxide scavenging (233.4± 14.2 μg/mL) etc. In addition, anti-inflammatory properties are also evident in terms of nitric oxide radical scavenging (28.76± 3.9 μg/mL) and lipoxygenase inhibition (39.2± 3.2 μg/mL) assays. Further, the anticancer activity of these extract has been identified against human endometrial and breast cancer cells. Further studies together with a bioassay guided fractionation may identify possible bioactive compounds.

**Highlights:**

- Thottea siliquosa is a traditional Ayurvedic medicine
- T. siliquosa exhibits significant anti-inflammatory and anticancer properties
- Polyphenolic compounds may be responsible for the bioactivities of T. siliquosa
- T. siliquosa may be a source for novel bioactive components

## 1. Introduction

Inflammation is the primary protective response of the body against exogenous molecules/ organisms. This involves a cascade of events which involve the recruitment of inflammatory cells to the site of action, release of inflammatory mediators from the granules of cells and subsequent tissue damage and edema. Acute inflammation is often described as protective response, as it doesn’t damage tissues for a long time. On the other hand, chronic inflammation, which lasts for a longer period of time are reported to be deleterious to the body.

Numerous studies have described the association of chronic inflammation with the onset and progression of various diseases such as cancers^1^, cardiovascular diseases^2^, fatty liver^3^, and Alzheimer’s disease^4^ Several studies have indicated that use of non-steroidal antiinflammatory drugs can reduce the incidence of various cancers^5^, cardiovascular diseases^6^, acute respiratory problems^7^, and Alzheimer’s disease^8^. Thus, inflammatory molecules have become the primary target for the prevention of various diseases.

Natural products has always been the source of drugs, likewise several different antiinflammatory drugs have been isolated from the Mother Nature^9,10^ In addition, the search for new anti-inflammatory drugs is also been intensified due to its multi-targeted nature and lack of side effects. Aristolochiaceae is an important family which is well known for its antioxidant and anticancer compound Aristolochicacids (AAs)^11^. The pharmacological properties of the family include anti-diarrhoeal^12^, anticancer^13^, anti-inflammatory and antipruritic activity^14^ Several novel alkaloid and terpenoids have also been isolated from the different plants belonging to this family^15^. The plants belonging to this family are also used traditionally in Chinese and Ayurvedic medicines as well as different folklore medicinal systems^16^. *Thottea siliquosa* is an unexplored medicinal plant which is traditionally used for digestive tract disorders and inflammatory conditions. Very limited literature is available on the biological and pharmacological properties of the plant. The only available published research indicates that the leaf and root extracts of *T. siliquosa* exhibits antibacterial activity against Streptococcus, E. coli and Pseudomonas strains^17^. Apart from these there are no available reports on the antioxidant and anti-inflammatory activities of the plant. Thus, the present study aimed to evaluate the antioxidant, anti-inflammatory and anticancer activities of the plant and further to elucidate the mechanism of action of the same.

## 2. Materials and Methods

### 2.1 Collection of plant material and extraction

The leaves of *Thottea siliquosa* plant was collected from Western Ghats regions of Kerala. The whole plant was cleaned properly, shade dried, and coarsely powdered. The powdered materials (10 g) were extracted with methanol (100 mL) by Soxhlet apparatus and the extracts obtained were concentrated under reduced pressure at 40°C using rotary evaporator. A portion of the residue was dissolved at 10 mg/mL stock solution in Dimethyl sulfoxide (DMSO) for the study and remaining portion was stored at 4°C for further use.

### 2.2 Cell lines and maintenance

Cells (HeLa and MCF7) were procured from National Centre for Cell Science, Pune and maintained at 5% carbon dioxide at 95% humidity. The cells were cultured in RPMI-1640 or DMEM media supplemented with 10% foetal bovine serum. The cells was passaged every third day or before reaching 80% confluency.

### 2.3 Total phenolic and total flavonoids content

Total phenolic content was quantified using a modified Folin-Ciocalteu method^18^. The results was compared to the standard curves and expressed as mg Gallic acid equivalent per gram dry powder for the samples.

Total flavonoid contents was quantified using the method explained by Khodaie, Bamdad, Delazar, Nazemiyeh^19^ and the results were expressed as mg quercetin equivalent per gram dry powder for the samples.

### 2.4 In vitro antioxidant activities

In vitro antioxidant capacity of the extract was determined in terms of DPPH radical scavenging^20^, Ferric reducing antioxidant power (FRAP)^21^, and hydrogen peroxide scavenging potentials.

### 2.5 In vitro anti-inflammatory activity

Lipoxygenase is an important mediator of inflammation, thereby making it a target for antiinflammatory activity. Inhibition of 5-Lipoxygenase was assessed according to the previously described methods^22^. Likewise nitric oxide scavenging activity had also been carried out as an indicator of anti-inflammatory potential. Percentage inhibition was calculated and IC_50_ was determined as per the methods described by Kakatum, Jaiarree, Makchucit, Itharat^23^.

### 2.6 Anti-hemolytic potential

The anti-hemolytic activity induced by a peroxyl radical initiator, AAPH was measured according to the method established by Salini, Divya, Chubicka, Meera, Fulzele, Ragavamenon, Babu^24^. Two hundred microliters of blood were mixed with 10 *μL* of plant extract at various concentrations (0-100 μg/mL) and then 600 *μ*L of AAPH (10%) was added. The mixture was incubated at 37° C for 30 minutes and the absorbance of the mixture was measured at 450 nm. The protective effect of the extract was expressed as the half maximum percentage inhibition of hemolysis induced by AAPH.

### 2.7 Anticancer activity

Cells were obtained from National Centre for Cell Sciences, Pune, India. They was maintained in RPMI-1640/ DMEM media, supplemented with 10% FBS at 37°C and 5% CO_2_. To study the anticancer activity, cancer cells (2.5×10^5^/ 250 *μL* volume) was plated in 48-well culture plates and was allowed to grow to subconfluency. Cells were then treated for 48 h with different concentrations of the plant extract (1 – 200 *μ*g/mL). The percentage cell viability of each was determined thereafter using determined using 3-(4,5-dimethyl thiazol-2-yl)-2, 5-diphenyl tetrazolium bromide-MTT assay^25^.

### 2.8 Statistical analysis

The values of in vitro studies were represented as mean± SD of four individual sets of the experiment, each conducted in triplicate. Statistical analysis was carried out by one way ANOVA followed by Tukey-Kramer multiple comparison tests using Graph Pad Prism software 7.0 versions.

## 3. Results

### 3.1 Total phenols and flavonoids content

The total phenol content of Methanolic extract of T. siliquosa was 4.67±0.41 GAE/ g of dry leaf. The flavonoid content of the extract was 1.88±0.22 QE/ g dry leaf (Figure 1).

**Figure 1.**
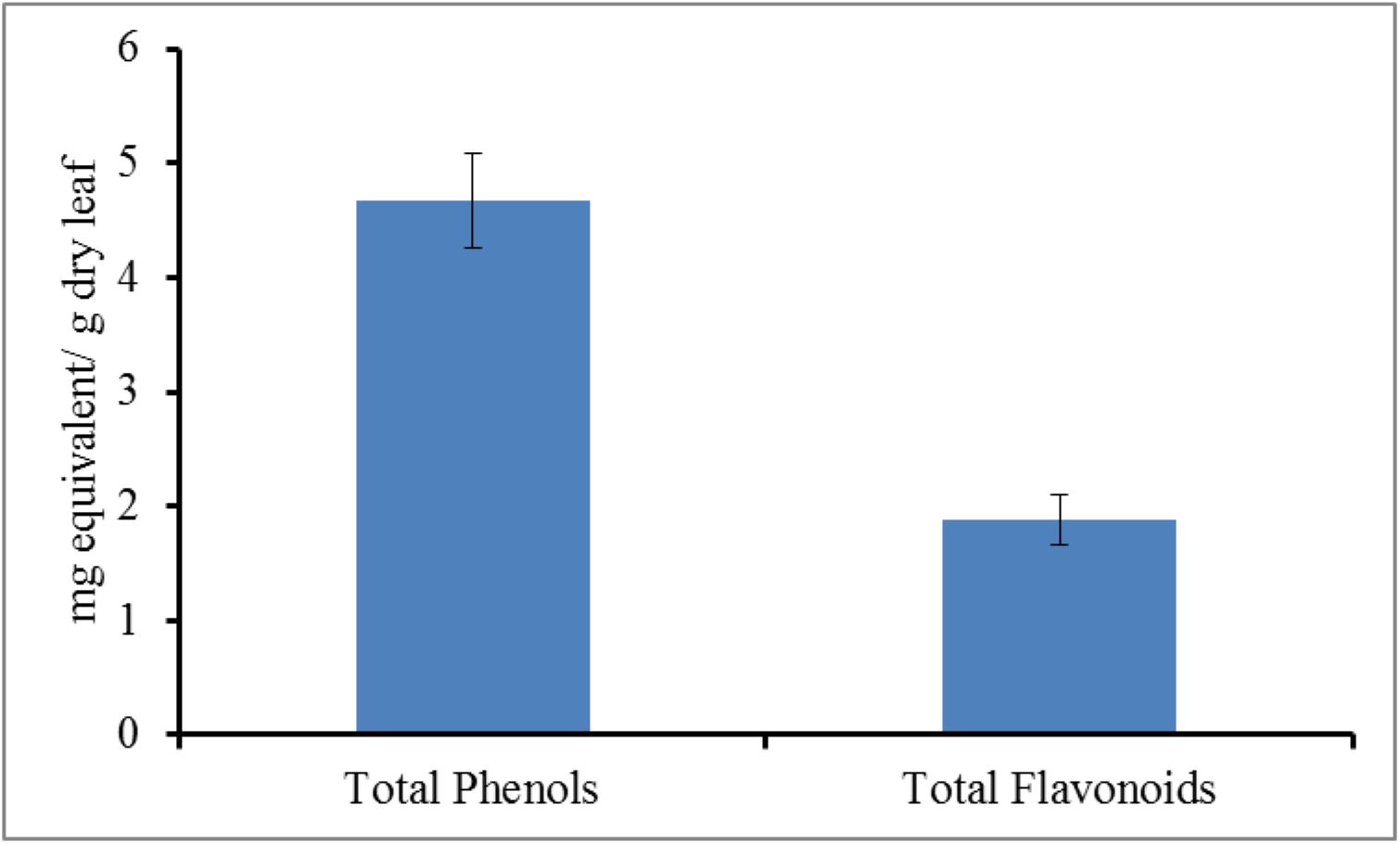
The total phenolic and total flavonoid contents in Thottea siliquosa Methanolic leaf extract.

### 3.2 In vitro antioxidant and anti-inflammatory potential

Antioxidant potential of the *Thottea siliquosa* methanolic extract has been evaluated in terms of DPPH radial scavenging, hydrogen peroxide radical scavenging, ferric reducing antioxidant power etc. The IC_50_ value of the extract in the DPPH scavenging was 110.40± 4.5; the half maximum inhibition value for hydrogen peroxide scavenging was 233.4± 14.2 μg/mL. The EC50 value in FRAP assay was found to be 41.1± 6.2 μg/mL (Table 1).

**Table 1.** Medium inhibition concentration for leaf methanolic extracts of *Thottea siliquosa* various in vitro antioxidant, anti-hemolytic and anti-inflammatory assays. For FRAP assay, EC50 concentration is given.

| Assay | IC<sub>50</sub> Value<br>(µg/mL) |
| --- | --- |
| DPPH radical scavenging | 110.40± 4.5 |
| Hydrogen peroxide scavenging | 233.4± 14.2 |
| Ferric reducing antioxidant power (EC<sub>50</sub>) | 41.1± 6.2 |
| Anti-hemolytic activity on RBC | 47.2±2.2 |
| Nitric oxide scavenging assay | 28.76± 3.9 |
| Lipoxygenase inhibition assay | 39.2± 3.2 |

The plant exhibited anti-inflammatory potential in lipoxygenase inhibition assay and nitric oxide scavenging. The IC_50_ values for the lipoxygenase inhibition and and nitric oxide scavenging assay were found to be 39.2± 3.2 and 28.76± 3.9 μg/mL (Table 1).

### 3.3 Anti-hemolytic potential and lipid peroxidation inhibition in RBC

As shown in Table 1, the IC_50_ values of hemolysis by T. siliquosa treatment were 47.2±2.2 *μ*g/mL. Morphological analysis also supported these results (Figure 2), where increased membrane damage caused by AAPH derived peroxyl radical was restored by the treatment with T. siliquosa extract. The lipid peroxidation in RBC was determined in terms of the TBARS, which was found to be 2.68±0.47 nmoles/mg Hb in untreated RBC, which was elevated to 7.12±0.62 nmoles/mg Hb in AAPH alone treated cells. TBARS level was reduced to 4.77±0.35 and 5.68±0.39 nmoles/mg Hb in cells pre-treated with T. siliquosa extracts (50 and 25 μg/mL).

**Figure 2.**
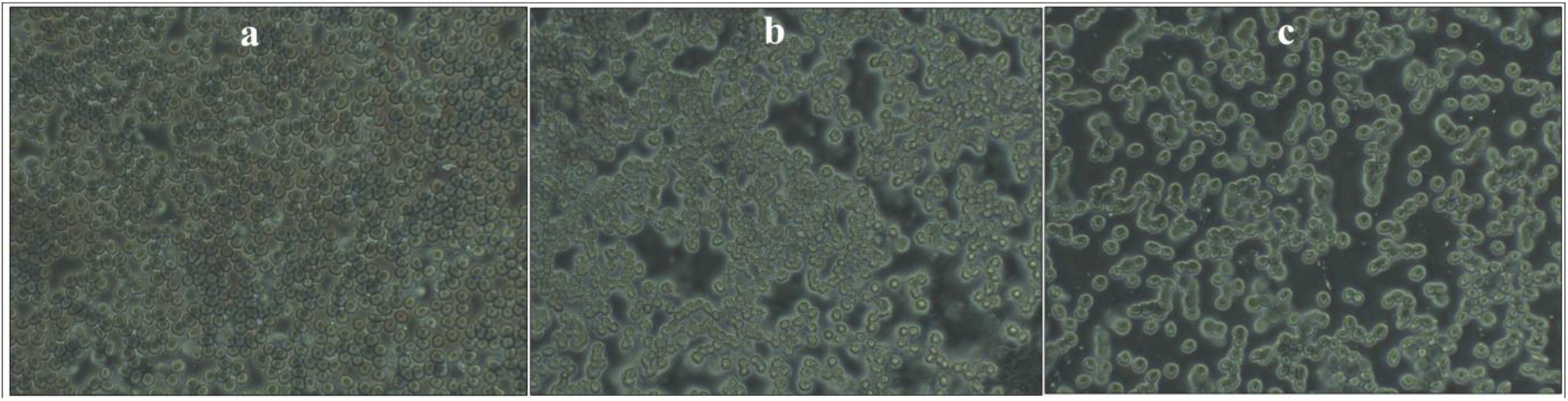
Morphological changes induced by AAPH treatment in RBCs and the protective effect by treatment with high dose of *T. siliquosa* extract. The normal (a), AAPH alone treated (b) and *T. siliquosa* extract treated (c) are shown.

### 3.4 In vitro anticancer activity

The anticancer potential of the extract was evaluated against HeLa (human endometrial cancer) and MCF7 (Human breast cancer) cells using MTT assay. The drug showed dose dependent decrease in the cell viability over a period of 48 hours and the IC_50_ value for the MCF7 cells was found to be 76.12± 5.5 μg/mL; the value was found to be 115.71± 10.2 μg/mL in HeLa cells (Figure 3).

**Figure 3.**
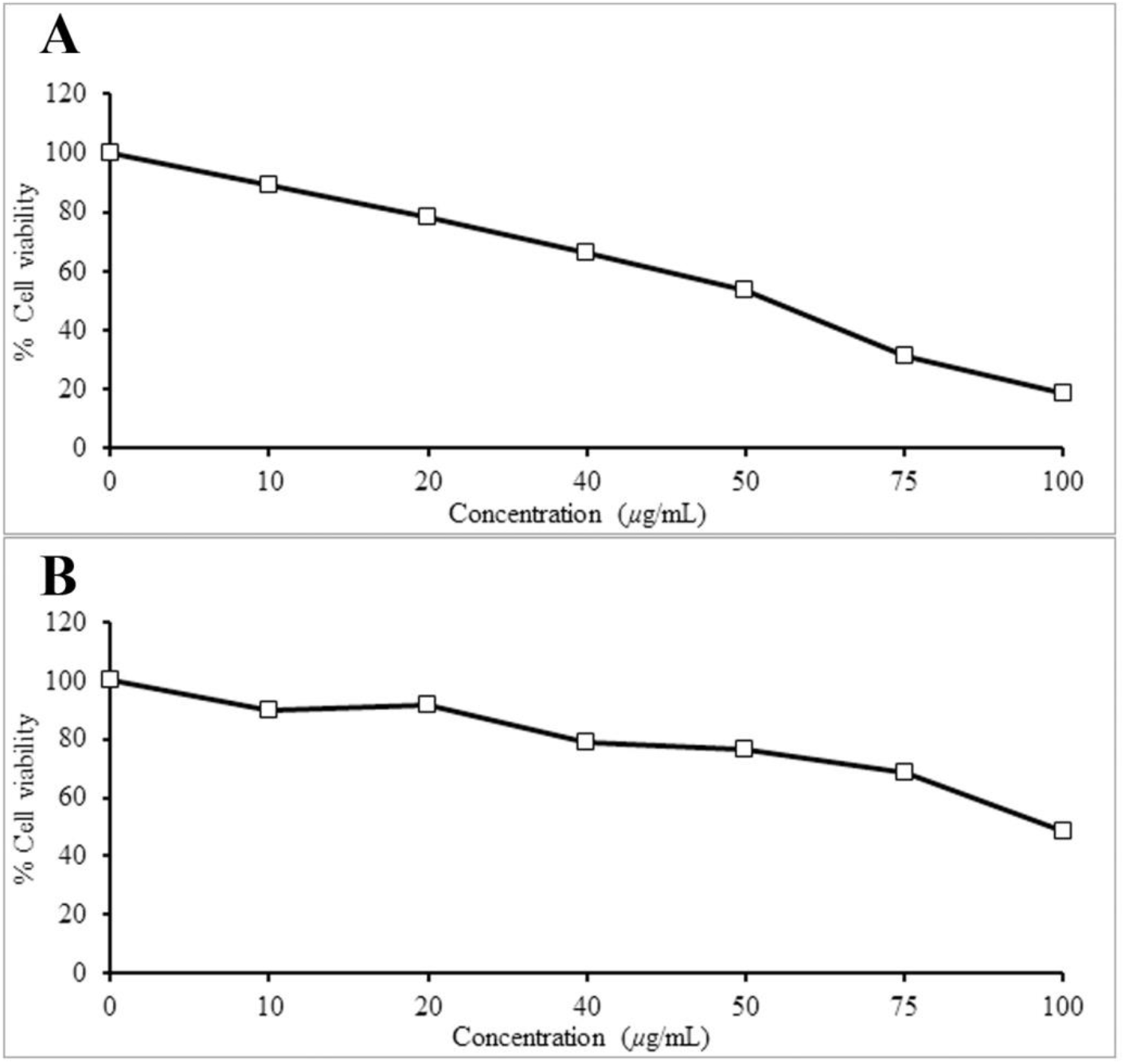
Anticancer potential of the T. siliquosa leaf methanolic extract against HeLa (a) MCF-7 cells (b).

## Discussion

Oxidative stress and inflammation are the two predominant driving forces behind various degenerative diseases. Hence, prevention of oxidative insults and chronic inflammation is the primary step for the prevention of these disorders. The present study analyzed the antioxidant and anti-inflammatory properties of a traditionally used Ayurvedic medicinal plant, Thottea siliquosa. Results show the moderate antioxidant properties of the plant in terms of DPPH and hydrogen peroxide radical scavenging properties and ferric reducing capacity. In addition to these, the extract also showed significant anti-inflammatory properties as indicated by the nitric oxide radical scavenging and lipoxygenase inhibition potential. By comparing the IC_50_ values for various antioxidant and anti-inflammatory assays, it can be assumed that the plant possess a high anti-inflammatory property than an antioxidant nature. The results also show that *T. siliquosa* efficiently reduces the damages induced by AAPH mediated oxidative toxicity, thereby improving the cell viability in human RBC.

Hydrogen peroxide and AAPH are significant sources of free radical generators, which are experimentally used to represent oxidative stress condition in cells. Among these, hydrogen peroxide derived radicals are the most potent in the induction of toxicity in mammalian cells.^26^. Azo compounds including AAPH are initiators of alkoxyl and peroxyl radicals in the hydrophilic compartments. Under physiological temperature, AAPH reduces automatically to form peroxyl radical (ROO*)^27,28^. Unlike the H_2_O_2_ radicals, alkoxyl and peroxyl radicals cannot pass through the cell membrane; whereas they react with the membrane leading to the peroxidation of membrane lipids^29^ and lipid peroxidation mediated oxidative DNA damages^30^. According to the reports by Sato, Kamo, Takahashi, Suzuki^31^, irrespective of the polar nature of AAPH, the generated peroxyl radicals diffuses through the lipid bilayer and be vested in in the nonpolar hydrocarbon interior. This increase the susceptibility of the cells to peroxidative changes, as hydrophobic region contains a considerably higher amounts of polyunsaturated fatty acids. This results in the initiation of a cascade of peroxidative events in the RBC membrane and thereby leading to hemolysis^32^. Corroborating with these reports, the present study also observed a significant reduction in the AAPH induced lipid peroxidation in RBC membrane. It is thus possible that the observed reduction in lipid peroxidation products in RBCs pre-treated with T. siliquosa may be the mechanism of antihemolytic properties of the plant.

Apart from these, the extract also showed significant anticancer properties against human endometrial and breast cancer cells. It is expected that the dose dependent anticancer properties may be mediated through apoptotic changes, however, further studies are necessary to ascertain these facts. The study also observed the presence of high amount of polyphenols and flavonoids in the T. siliquosa extract, which might be responsible for the protective effects observed against free radicals and anti-inflammatory potential. In addition, these bioactive polyphenol compounds are also known for their anticancer potentials^33–36^. It is therefore expected that further purification of the extract may be helpful to identify novel bioactive compounds with antioxidant, anti-inflammatory and anticancer properties.

## Acknowledgement

The authors acknowledge Rashtriya Uchchatar Shiksha Abhiyan (A1/SJC/RUSA-SPD/MRP/60/2019) financial support in the form of minor research project and Kerala Science Council for Science, Technology and Environment Student Project Scheme.

## Reference

1. Munn LL. Cancer and inflammation. Wiley Interdisciplinary Reviews: Systems Biology and Medicine. 2017;9(2):e1370.

2. Zhong X, Guo L, Zhang L, Li Y, He R, Cheng G. Inflammatory potential of diet and risk of cardiovascular disease or mortality: A meta-analysis. Scientific Reports. 2017;7(1):6367.

3. Del Campo JA, Gallego P, Grande L. Role of inflammatory response in liver diseases: Therapeutic strategies. World Journal of Hepatology. 2018;10(1):1–7.

4. Whittington RA, Planel E, Terrando N. Impaired Resolution of Inflammation in Alzheimer’s Disease: A Review. Frontiers in Immunology. 2017;8(1464).

5. Doat S, Cénée S, Trétarre B, et al. Nonsteroidal anti-inflammatory drugs (NSAIDs) and prostate cancer risk: results from the EPICAP study. Cancer Medicine. 2017;6(10):2461–2470.

6. Dolgin E. Anti-inflammatory drug cuts risk of heart disease — and cancer. Nature Reviews Drug Discovery. 2017;16:665.

7. Wen YC, Hsiao FY, Chan KA, Lin ZF, Shen LJ, Fang CC. Acute Respiratory Infection and Use of Nonsteroidal Anti-Inflammatory Drugs on Risk of Acute Myocardial Infarction: A Nationwide Case-Crossover Study. J Infect Dis. 2017;215(4):503–509.

8. Ardura-Fabregat A, Boddeke EWGM, Boza-Serrano A, et al. Targeting Neuroinflammation to Treat Alzheimer’s Disease. CNS Drugs. 2017;31(12):1057–1082.

9. Arulselvan P, Fard MT, Tan WS, et al. Role of Antioxidants and Natural Products in Inflammation. Oxidative Medicine and Cellular Longevity. 2016;2016:5276130.

10. Orlikova B, Legrand N, Panning J, Dicato M, Diederich M. Anti-Inflammatory and Anticancer Drugs from Nature. Paper presented at: Advances in Nutrition and Cancer; 2014//, 2014; Berlin, Heidelberg.

11. Kuo P-C, Li Y-C, Wu T-S. Chemical Constituents and Pharmacology of the Aristolochia ( mădōu ling) species. Journal of traditional and complementary medicine. 2012;2(4):249–266.

12. Dharmalingam SR, Madhappan R, Ramamurthy S, et al. Investigation on antidiarrhoeal activity of Aristolochia indica Linn. Root extracts in mice. African journal of traditional, complementary, and alternative medicines: AJTCAM. 2014;11(2):292–294.

13. Hadem KLH, Sharan RN, Kma L. Phytochemicals of Aristolochia tagala and Curcuma caesia exert anticancer effect by tumor necrosis factor-α-mediated decrease in nuclear factor kappaB binding activity. Journal of basic and clinical pharmacy. 2015;7(1):1–11.

14. Mathew JE, Kaitheri SK, Dinakaranvachala S, Jose M. Anti-inflammatory, Antipruritic and Mast Cell Stabilizing Activity of Aristolochia Indica. Iran J Basic MedSci. 2011;14(5):422–427.

15. Chung YM, Chang FR, Tseng TF, et al. A novel alkaloid, aristopyridinone A and anti-inflammatory phenanthrenes isolated from Aristolochia manshuriensis. Bioorg Med Chem Lett. 2011;21(6):1792–1794.

16. Wu TS, Damu AG, Su CR, Kuo PC. Terpenoids of Aristolochia and their biological activities. Nat Prod Rep. 2004;21(5):594–624.

17. Nusaiba SAW, Murugan K. In vitro analysis on bactericidal screening and antioxidant potentiality of leaf and root extracts of Thottea siliquosa (Lam.) Ding Hou. An ethnobotanical plant. Asian Pacific Journal of Tropical Biomedicine. 2013;3(11):859–865.

18. Illam SP, Narayanankutty A, Raghavamenon AC. Polyphenols of virgin coconut oil prevent pro-oxidant mediated cell death. Toxicology mechanisms and methods. 2017;27(6):442–450.

19. Khodaie L, Bamdad S, Delazar A, Nazemiyeh H. Antioxidant, total phenol and flavonoid contents of two pedicularis L. Species from eastern azerbaijan, iran. Bioimpacts. 2012;2(1):43–57.

20. Lai SC, Ho YL, Huang SC, et al. Antioxidant and antiproliferative activities of Desmodium triflorum (L.) DC. Am J Chin Med. 2010;38(2):329–342.

21. Konate K, Souza A, Coulibaly AY, et al. In vitro antioxidant, lipoxygenase and xanthine oxidase inhibitory activities of fractions from Cienfuegosia digitata Cav., Sida alba L. and Sida acuta Burn f (Malvaceae). Pak J Biol Sci. 2010;13(22):1092–1098.

22. Ben-Nasr S, Aazza S, Mnif W, Miguel MdGC. Antioxidant and anti-lipoxygenase activities of extracts from different parts of Lavatera cretica L. grown in Algarve (Portugal). Pharmacognosy Magazine. 2015;11(41):48–54.

23. Kakatum N, Jaiarree N, Makchucit S, Itharat A. Antioxidant and anti-inflammatory activities of Thai medicinal plants in Sahasthara remedy for muscle pain treatment. J Med Assoc Thai. 2012;95(1):S120–126.

24. Salini S, Divya MK, Chubicka T, et al. Protective effect of Scutellaria species on AAPH-induced oxidative damage in human erythrocyte. Journal of basic and clinical physiology and pharmacology. 2015;15(10):2015–0081.

25. Mosmann T. Rapid colorimetric assay for cellular growth and survival: application to proliferation and cytotoxicity assays. J Immunol Methods. 1983;65(1-2):55–63.

26. Nakamura J, Purvis ER, Swenberg JA. Micromolar concentrations of hydrogen peroxide induce oxidative DNA lesions more efficiently than millimolar concentrations in mammalian cells. Nucleic Acids Res. 2003;31(6):1790–1795.

27. Werber J, Wang YJ, Milligan M, Li X, Ji JA. Analysis of 2,2’-azobis (2-amidinopropane) dihydrochloride degradation and hydrolysis in aqueous solutions. J Pharm Sci. 2011;100(8):3307–3315.

28. Niki E. Free radical initiators as source of water-or lipid-soluble peroxyl radicals. Methods Enzymol. 1990; 186:100–108.

29. Ayala A, Munoz MF, Arguelles S. Lipid Peroxidation: Production, Metabolism, and Signaling Mechanisms of Malondialdehyde and 4-Hydroxy-2-Nonenal. Oxidative Medicine and Cellular Longevity. 2014;2014:31.

30. Lim P, Wuenschell GE, Holland V, et al. Peroxyl radical mediated oxidative DNA base damage: implications for lipid peroxidation induced mutagenesis. Biochemistry. 2004;43(49):15339–15348.

31. Sato Y, Kamo S, Takahashi T, Suzuki Y. Mechanism of Free Radical-Induced Hemolysis of Human Erythrocytes: Hemolysis by Water-Soluble Radical Initiator. Biochemistry. 1995;34(28):8940–8949.

32. Banerjee A, Kunwar A, Mishra B, Priyadarsini KI. Concentration dependent antioxidant/pro-oxidant activity of curcumin studies from AAPH induced hemolysis of RBCs. Chemico-biological interactions. 2008;174(2):134–139.

33. Narayanankutty A. Toll-like Receptors as a Novel Therapeutic Target for Natural Products Against Chronic Diseases. Curr Drug Targets. 2019;20(10):1068–1080.

34. Narayanankutty A. PI3K/ Akt/ mTOR Pathway as a Therapeutic Target for Colorectal Cancer: A Review of Preclinical and Clinical Evidence. Curr Drug Targets. 2019;20(12):1217–1226.

35. Narayanankutty V, Narayanankutty A, Nair A. Heat Shock Proteins (HSPs): A Novel Target for Cancer Metastasis Prevention. Curr Drug Targets. 2019;20(7):727–737.

36. Shweta M, Arunaksharan N. Traditional Fruits of Kerala: Bioactive Compounds and their Curative Potential in Chronic Diseases. Current Nutrition & Food Science. 2017;13(4):279–289.

